# Delving into human α1,4-galactosyltransferase acceptor specificity: the role of enzyme dimerization

**DOI:** 10.1101/2024.03.21.586141

**Authors:** Krzysztof Mikołajczyk, Karol Wróblewski, Sebastian Kmiecik

## Abstract

Human α1,4-galactosyltransferase (A4galt), a Golgi apparatus-resident GT, synthesizes Gb3 glycosphingolipid (GSL) and P1 glycotope on glycoproteins (GPs), which are receptors for Shiga toxin types 1 and 2. Despite the significant role of A4galt in glycosylation processes, the molecular mechanisms underlying its varied acceptor specificities remain poorly understood. Here, we attempted to elucidate A4galt specificity towards GSLs and GPs by exploring its interaction with GTs with various acceptor specificities, GP-specific β1,4-galactosyltransferase 1 (B4galt1) and GSL-specific β1,4-galactosyltransferase isoenzymes 5 and 6 (B4galt5 and B4galt6). Using a novel NanoBiT assay, we found that A4galt can form homodimers and heterodimers with B4galt1 and B4galt5 in two cell lines, human embryonic kidney cells (HEK293) and Chinese hamster ovary cells (CHO-Lec2). We found that A4galt-B4galts heterodimers preferred N-terminally tagged interactions, while in A4galt homodimers, the favored localization of the fused tag depended on the cell line used. Furthermore, by employing AlphaFold for state-of-the-art structural prediction, we analyzed the interactions and structures of these enzyme complexes. Our analysis highlighted that the A4galt-B4galt5 heterodimer exhibited the highest prediction confidence, indicating a significant role of A4galt heterodimerization in determining enzyme specificity toward GSLs and GPs. These findings enhance our knowledge of A4galt acceptor specificity and may contribute to a better comprehension of pathomechanisms of the Shiga toxin-related diseases.

## Introduction

Human α1,4-galactosyltransferase (A4galt, Gb3/CD77 synthase, P1/P^k^ synthase, EC 2.4.1.228) is a type II transmembrane glycosyltransferase (GT) located in the *trans*-Golgi apparatus. This enzyme consists of a C-terminal globular catalytic domain oriented towards the Golgi lumen and an N-terminal cytoplasmic domain [1–2]. Topologically, human A4galt comprises a cytoplasmic domain (spanning 1-22 amino acid residues), a transmembrane domain (23-43 aa residues), which anchors the enzyme within the Golgi membrane, and a luminal domain (44-353 aa residues), containing the catalytic site (according to UniProt Q9NPC4, https://www.uniprot.org/).

*In silico* analysis of the human A4galt AlphaFold model revealed that the enzyme belongs to the GT-A fold-type GT and is a member of 32 families in the CAZy glycosyltransferase database (Carbohydrate Active Enzymes Database, CAZy, http://www.cazy.org/). For catalytic activity, the enzyme requires a divalent metal ion (typically Mn^2+^) coordinated by two aspartic acid residues to form a DXD motif (D_192_TD in human A4galt) [1,3]. Human A4galt occurs as a high-frequency enzyme, containing a glutamine (Q) at position 211 (p.Q211), and a rare enzyme variant (referred to as mutein) with a p.Q211E substitution (rs397514502), found in only two families worldwide [4]. This amino acid substitution affects the enzyme acceptor specificity towards GalNAc-capped oligosaccharides, enabling the synthesis of NOR1 and NOR2 antigens (terminally with a Galα1→4GalNAc disaccharide) [2,4–5].

For many years, human A4galt has been considered as a glycosphingolipid (GSL)-specific enzyme, responsible for transferring Gal residues into GSL acceptors, such as lactosylceramide, resulting in the formation of globotriaosylceramide (Gb3, Galα1→4Galβ1→4Glc-Cer) [2,5]. However, recent studies have shown that A4galt reveals a broader enzyme specificity toward glycoprotein (GP)-related acceptors, particularly N-glycans. It may produce a unique oligosaccharide motif, named P1 glycotope (consisting of a Galα1→4Galβ1→4GlcNAc carbohydrate sequence), which contains a Galα1→4Galβ disaccharide, identical to the terminal disaccharide of Gb3 [6]. Both products generated by A4galt, Gb3 on GSLs and the P1 glycotope on glycoproteins, may serve as receptors for Shiga toxins (Stxs) [6–8]. Shiga toxin-producing *Escherichia coli* (STEC) strains, similar to *Shigella dysenteriae* serotype 1, are posing an increasing threat to the human population. STEC infections lead to hemorrhagic colitis, which often progresses to hemolytic-uremic syndrome (HUS), a severe complication that usually causes acute kidney failure [9–11]. Stxs are produced by STEC are of two types: Stx1, which is identical to the toxin secreted by *S. dysenteriae* serotype 1, and the more distinct Stx2. Structurally, Stxs belong to the AB_5_ toxin family and comprise an A catalytic subunit responsible for cytotoxic effects and a pentameric B subunit that specifically binds to cell receptors (Gb3 and P1 glycotope) [6,9,12–13]. Stx1 exhibits a remarkable ability to utilize not only the GSL receptor but also the P1 glycotope on N-glycoproteins for cell entry and cytotoxicity, whereas Stx2 exclusively relies on GSL-related receptors [6,8].

Herein, we propose that human A4galt forms complexes within the Golgi apparatus, orchestrating sequential glycan biosynthesis pathways that utilize GSL- and GP-based acceptors [14]. Preliminary data have indicated that human A4galt forms homodimers [15] and heterodimers with β1,4-galactosyltransferase 6 (B4galt6) [16]. To investigate the dimer formation of human A4galt, we used a relatively new NanoBiT (NanoLuc® Binary Technology) Protein: Protein Interaction System in CHO-Lec2 and HEK293T cells. To further understand the molecular mechanisms underlying A4galt homo- and heterodimerization, we also used computational tools based on the AlphaFold model of the enzymes.

## Material and methods

### Cell cultures

CHO-Lec2 and HEK293T cells were obtained from the American Type Culture Collection (Rockville, MD, USA). The cells were grown and maintained in a humidified incubator with 5% CO2 at 37°C in DMEM/F12 medium (Thermo Fisher Scientific, Inc., Waltham, MA, USA) supplemented with 10% fetal bovine serum (Gibco, Inc., Waltham, MA, USA) and Pen-Strep (Gibco, Inc., Waltham, MA, USA). The culture medium was changed every second or third day, and after reaching 85–90% confluence, the cells were subcultured by treatment with trypsin (0.25% trypsin, 137 mM NaCl, 4.3 mM NaHCO3, 5.4 mM KCl, 5.6 mM glucose, 0.014 mM phenol red, 0.7 mM EDTA), harvested, centrifuged at 800 × g for 5 min, resuspended in fresh medium and seeded to new tissue culture plates.

### Construction of expression vectors

The ORF sequences of human *B4GALT1* (GenBank accession number NM_001378495.1: 30-1187), *B4GALT5* (GenBank accession number NM_004776.4: 189-1355), and *B4GALT6* (GenBank accession number NM_004775.5: 156-1187) were customized by Thermo Fisher Scientific (Inc., Waltham, MA, USA) and cloned into pET-TOPO plasmids (Table SI). For human *A4GALT*, the template for PCR amplification was the pCAG-A4GALT expression plasmid containing the full-length human *A4GALT* gene (GenBank accession number NM_001318038.3: nucleotides 325-1386), which includes both the high-frequency enzyme variant and the c.631C>G substitution variant (rs397514502) (Table SI). The expression constructs for the split luciferase complementation assay (pBiT1.1-C [TK/LgBiT], pBiT2.1-C [TK/SmBiT], pBiT1.1-N [TK/LgBiT] and pBiT2.1-N [TK/SmBiT]) were prepared according to the manufacturer’s protocol (NanoLuc® Binary Technology NanoBiT™, Promega, Madison, WI, USA). Human *B4GALT1*, *B4GALT5*, and *B4GALT6* genes, along with human *A4GALT* (high-frequency and mutein form), were cloned into pBiT1.1-N/C and pBiT2.1-N/C vectors. These vectors encoded LgBiT and SmBiT luciferase subunits at the N- or C-terminus of the protein interaction candidates (presented in Fig. SI). PCR was performed using an MJ Mini gradient PCR apparatus (BioRad, Hercules, CA, USA) in 20 µl reaction mixes containing 200 ng of genomic DNA, 0.2 mM dNTPs, Taq buffer with KCl (1:10 dilution), 1.5 mM MgCl_2_, 0.2 mM forward and reverse primers, and 1 unit of Taq polymerase (Thermo Fisher Scientific, Inc., Waltham, MA, USA). The PCR conditions are shown in Table SII. Cloning of pBiT vectors was performed using restriction sites added to the forward and reverse primers, according to the primers listed in Table SIII. Ligation of inserts into the pBiT vectors was carried out using T4 DNA Ligase (Promega, Madison, WI, USA). To obtain pBiT constructs, *Escherichia coli* DH5α (ElectroMAX™ DH5α-E Competent Cells, Thermo Fisher Scientific, Inc., Waltham, MA, USA) were transformed with proper expression constructs by electroporation. Positive clones were selected and used for large-scale plasmid production using *E. coli* DH5α. Plasmids were subsequently isolated using the GeneJET Plasmid Miniprep Kit (Thermo Fisher Scientific, Inc., Waltham, MA, USA) and sequenced as described in [1,6].

### Split luciferase complementation assay

CHO-Lec2 and HEK293T cells (2×10^4^) were seeded in a complete growth medium in a 96-well plate with white polystyrene wells and a flat transparent bottom (Corning Inc., New York, USA). After 20–24 h, the cells were transfected with appropriate combinations of plasmids (25 ng/well of each plasmid, 50 ng/well in total) using a linear polyethylenimine (PEI) 25 kDa (1 mg/mL, Polysciences Inc.) transfection reagent (DNA:PEI ratio 1:2). Approximately 2–3 h before the measurement, the conditioned medium was replaced with 100 µl of serum-free Opti-MEM medium (Life Technologies, CA, USA). Immediately before measurement, Nano-Glo® Live Cell Substrate (Promega) was added to all wells, according to the manufacturer’s instructions. Cell-derived luminescence was recorded using a Clariostar Luminescence Microplate Reader (BMG LABTECH). Each tested combination was accompanied by the respective negative control comprising the analyzed GT fused with the large NanoLuc subunit combined with the HaloTag (a recombinant protein not synthesized by mammalian cells, thus not interacting with proteins of mammalian origin) that was fused with the small NanoLuc subunit (Promega).

### Data analysis

For statistical analysis, one-way ANOVA with Bonferroni post-hoc test was used. All analyses were performed with GraphPad Prism (GraphPad Software, CA, USA). Statistical significance was assigned to *p*-value < 0.05. Only results exceeding the respective negative controls by at least 10-fold were considered indicative of an interaction, as suggested by the manufacturer.

### Dimer structure prediction

Structure prediction was carried out using AlphaFold-Multimer [17–18] implemented in ColabFold [19]. Multiple parameters were tested, such as template modes, templates used, dropout, and the number of recycles. These tests yielded an array of structure predictions. The final dimer models for structural analysis were selected using two criteria: (1) the lowest average PAE score for the residue pairs that were in contact (< 6 Å apart) and (2) the highest average pLDDT score.

### Dimer structure analysis

Structure Prediction and Omics-based Classifier (SPOC) is a machine learning tool that integrates structural predictions with omics data to assess the reliability of predicted protein-protein interactions [20]. It is valuable for distinguishing true interactions from false positives, offering a robust metric that enhances confidence in computational biology studies. This tool takes 3 user-uploaded AlphaFold multimer (AF-M) predictions and calculates compact SPOC scores and other confidence metrics (avg_models, ipTM, pDOCKQ). For each dimer, the three highest scored models of the five models generated by AlphaFold-Multimer were used for this analysis.

### Coevolution analysis

The coevolution analysis was performed using EVcouplings [21]. The method generates multiple sequence alignments (MSA) for each of the two sequences using Jackhammer (with a Bitscore cutoff of 0.3) and then combines them into a single combined MSA. The measure of whether MSA is sufficiently deep is the number of effective sequences divided by the length of the protein, which should be above 1.

### Active site prediction

The potential active sites were predicted using the COACH meta-server (https://zhanggroup.org/COACH/) [22]. It is a consensus method that uses multiple methods to predict the potential active sites. These methods include: COACH, TM-SITE, S-SITE, COFACTOR, FINDSITE and ConCavity. Only methods that correctly predicted the known substrates for these proteins were considered. The residues were considered as a part of the active site if the majority of methods predicted them to form the active center.

### Visualization

The following tools were used for visualization figure preparation: PyMol [The PyMOL Molecular Graphics System, Version 3.0 Schrödinger, LLC.] (Fig. 3 and Fig. SIII-SVI), Mapiya web-server [23] (Fig. 4-5), PAE Viewer [24] (Fig. SII).

## Results

### Analysis of human A4galt dimerization using NanoBiT technology

The specificity towards GP- and GSL-based acceptors of human A4galt may be influenced by its interaction with other GTs, which vary in acceptor specificities. As potential dimerization partners for human A4galt, we contemplated representatives of β1,4-galactosyltransferases (B4galt1-4 and B4galt5-6). B4galt1 is a GT located in the *trans*-Golgi that synthesizes β-galactosylated N-glycans (GP-based substrate for human A4galt), whereas B4galt2-4 also exhibits glycoprotein-utilizing activity, albeit to a lesser extent [25–26]. B4galt5 and B4galt6 catalyze the synthesis of lactosylceramide (LacCer), a major GSL-based substrate for human A4galt. In addition, both enzymes are located in the Golgi cisternae and the *trans*-Golgi network (TGN) [27]. The co-occurrence of these enzymes in *trans*-Golgi and involvement in the synthesis of A4galt substrates (Gal-terminated N-glycans and LacCer) suggests that heterodimers containing A4galt and B4galt1-4 and/or B4galt5-6 may exist. Their spatial proximity in glycosylation pathways increases the likelihood of interactions, potentially contributing to the formation of supramolecular complexes involving these GTs. To analyze heterodimer assembly, we selected B4galt1, a major isoenzyme of GPs-specific β1,4-galactosyltransferases [28], and B4galt5 and B4galt6, two major GSLs-specific isoenzymes [29–30]. The presence of such heterocomplexes could provide insight into the dual specificity of human A4galt towards Gal-terminated GSL- and GP-based acceptors.

NanoBiT technology was used to assess the formation of homo- and heterodimers by high-frequency and mutein human A4galt. Although it is a relatively new method, NanoBiT has already been used to identify interaction partners for other GTs, such as human B4galt4 [31] and B4galt1 [32], as well as other proteins [33–34]. In our study, we used two cell lines, CHO-Lec2 and HEK293T. Previously, CHO-Lec2 cells were transfected with human *A4GALT* and used to evaluate the ability of human A4galt to recognize GSL- and GP-based acceptors [6, 35]. Since CHO-Lec2 cells do not contain the homolog of the *A4GALT* gene, which encodes A4galt, these cells are not primarily sensitive to Stxs. Additionally, these cells are deprived of CMP-sialic acid transporter, thus cannot produce sialic acid-terminated N-glycans, resulting in the dominance of galactosylated oligosaccharides on the cell surface (allowing a better Stxs affinity to cells) [6]. To evaluate whether the results obtained from CHO-Lec2 cells corresponded to the human cell line, we conducted the same NanoBiT experiments using HEK293T cells which express human A4galt and can bind Stxs.

Using NanoBiT, we tested various combinations of proteins fused with the LgBiT and SmBiT subunits (located on either the N- or C-terminus of the analyzed protein) to determine the optimal orientation for interactions (Table SIII). We found that both forms of human A4galt (high-frequency and mutein) formed homodimers in both CHO-Lec2 and HEK293T cells (Fig. 1 and 2). Notably, the highest average luminescence value (compared to the respective controls) was observed for mutein in CHO-Lec2 cells (Fig. 2), in contrast to HEK293T cells (Fig. 1), in which the average luminescence of A4galt homodimers was similar for both high-frequency and mutein forms. We also found that N-terminally tagged constructs were preferred for heterodimer formation by human A4galt, while in the case of homodimers, the favored localization of the fused tag depended on the cell line used. In HEK293T cells, a high-frequency enzyme preference tag occurs at the C-terminus for homodimerization (Fig. 1A, B), while the mutein enzyme favored the N-terminally tagged constructs (Fig. 1C, D). High-frequency human A4galt in CHO-Lec2 cells formed homodimers using both N- and C-tagged constructs (Fig. 2A, B). The roles of A4galt N- and C-termini in PPIs of different cellular types require further investigation.

**Fig. 1.**
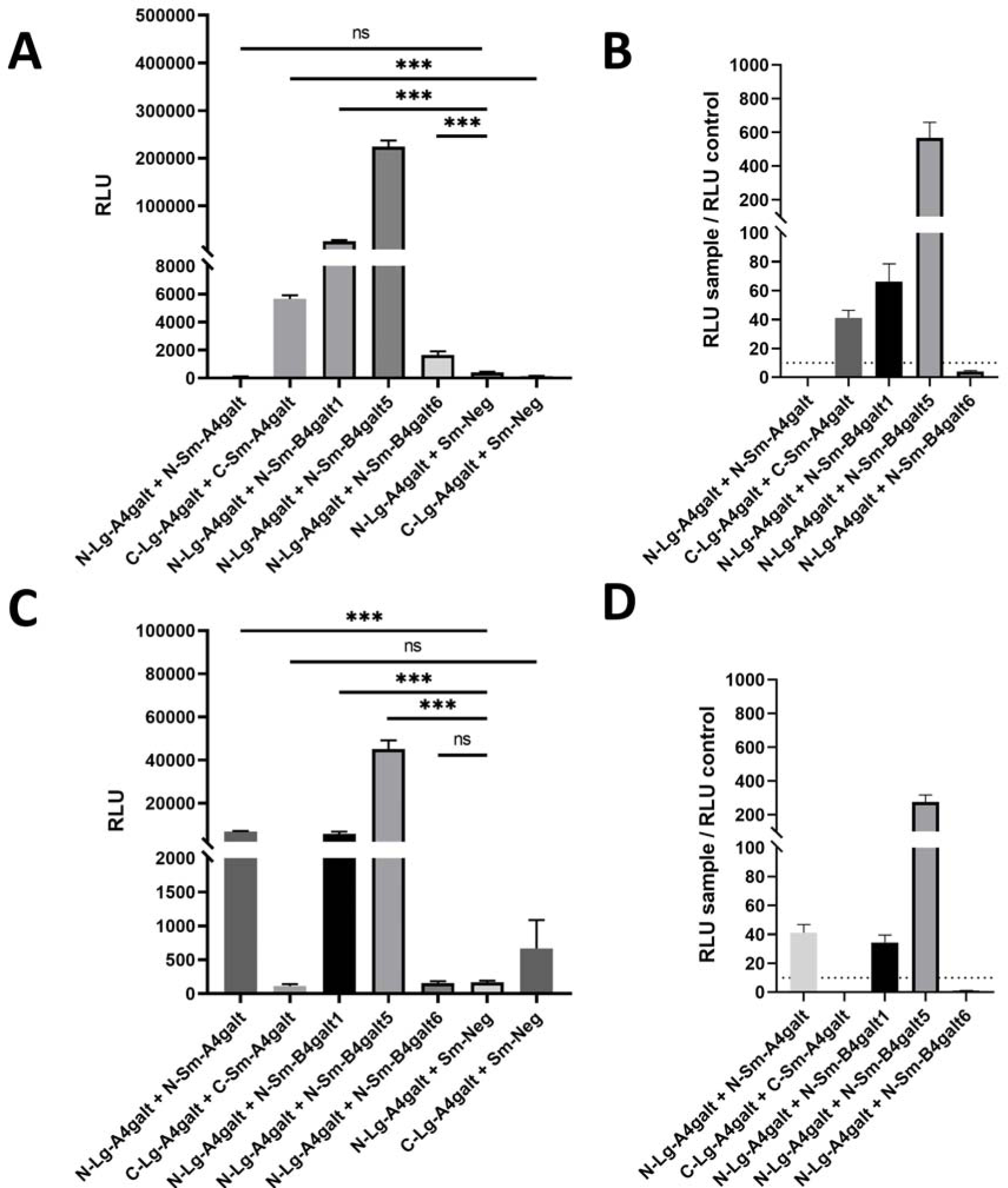
NanoBiT assay was performed for high-frequency human A4galt (**A**) and its mutein form (**C**) with the potential heterologous protein partners (B4galt1, B4galt5, and B4galt6 isoenzyme) using HEK293T cells. The negative controls comprise HaloTag fused with the small NanoLuc subunit. The fold changes of luminescence were calculated by dividing the average luminescence measured for the tested combinations (Sample RLU) was divided by the average luminescence obtained for the corresponding negative controls (Control RLU) to calculate the fold changes of the interaction pairs for high-frequency human A4galt (**B**) and its mutein (**D**). The threshold value was set at 10, according to the manufacturer. N-Lg-A4galt, human A4galt N-terminated tag with the LgBiT NanoLuc luciferase subunit; N-Lg-A4galt/B4galt1/B4galt5/B4galt6, human α1,4-galactosyltransferase/β1,4-galactosyltransferase 1/β1,4-galactosyltransferase 5 isoenzyme/β1,4-galactosyltransferase 6 isoenzyme N-terminated fused with the LgBiT NanoLuc luciferase subunit; C-Lg-A4galt/B4galt1/B4galt5/B4galt6, human α1,4-galactosyltransferase/β1,4-galactosyltransferase 1/β1,4-galactosyltransferase 5 isoenzyme/β1,4-galactosyltransferase 6 isoenzyme C-terminated fused with the LgBiT NanoLuc luciferase subunit; N-Sm-A4galt/B4galt1/B4galt5/B4galt6, human α1,4-galactosyltransferase/β1,4-galactosyltransferase 1/β1,4-galactosyltransferase 5 isoenzyme/β1,4-galactosyltransferase 6 isoenzyme N-terminated fused with the SmBiT NanoLuc luciferase subunit; C-Sm-A4galt/B4galt1/B4galt5/B4galt6, human α1,4-galactosyltransferase/β1,4-galactosyltransferase 1/β1,4-galactosyltransferase 5 isoenzyme/β1,4-galactosyltransferase 6 isoenzyme C-terminated fused with the SmBiT NanoLuc luciferase subunit. Statistical significance was established as ∗, p < 0.05; ∗∗p < 0.01; ∗∗∗p < 0.001; and ∗∗∗∗p < 0.0001; ns, not significant.

**Fig. 2.**
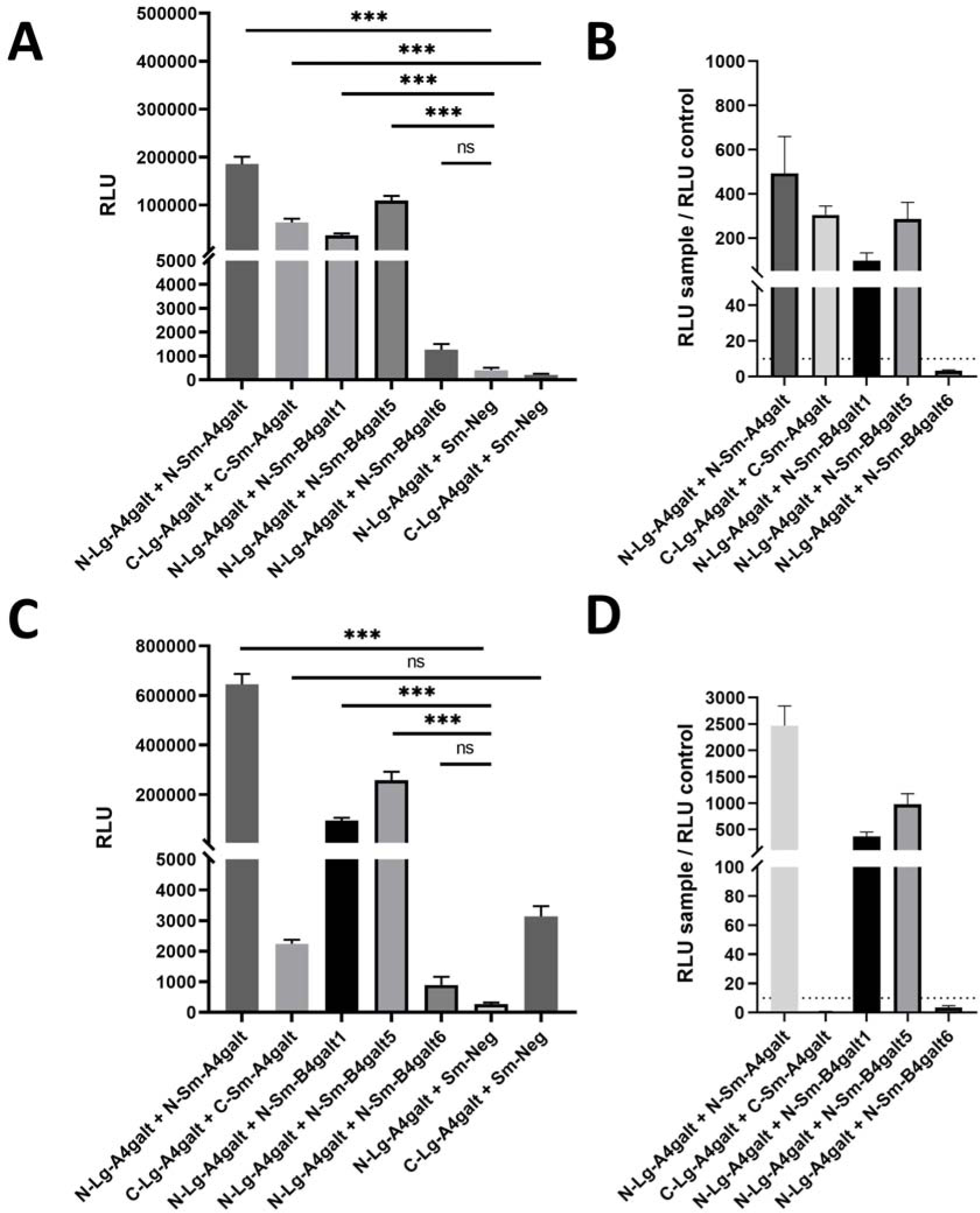
NanoBiT assay performed for high-frequency human A4galt (**A**) and its mutein form (**C**) with the potential heterologous protein partners (B4galt1, B4galt5, and B4galt6 isoenzyme) using CHO-Lec2 cells. The negative controls comprise HaloTag fused with the small NanoLuc subunit. The fold changes of luminescence were calculated by dividing the average luminescence measured for the tested combinations (Sample RLU) was divided by the average luminescence obtained for the corresponding negative controls (Control RLU) to calculate the fold changes of the interaction pairs for high-frequency human A4galt (**B**) and its mutein (**D**). The threshold value was set at 10, according to the manufacturer. N-Lg-A4galt, human α1,4-galactosyltransferase N-terminated tag with the LgBiT NanoLuc luciferase subunit; N-Lg-A4galt/B4galt1/B4galt5/B4galt6, human α1,4-galactosyltransferase/β1,4-galactosyltransferase 1/β1,4-galactosyltransferase 5 isoenzyme/β1,4-galactosyltransferase 6 isoenzyme N-terminated fused with the LgBiT NanoLuc luciferase subunit; C-Lg-A4galt/B4galt1/B4galt5/B4galt6, human α1,4-galactosyltransferase/β1,4-galactosyltransferase 1/β1,4-galactosyltransferase 5 isoenzyme/β1,4-galactosyltransferase 6 isoenzyme C-terminated fused with the LgBiT NanoLuc luciferase subunit; N-Sm-A4galt/B4galt1/B4galt5/B4galt6, human α1,4-galactosyltransferase/β1,4-galactosyltransferase 1/β1,4-galactosyltransferase 5 isoenzyme/β1,4-galactosyltransferase 6 isoenzyme N-terminated fused with the SmBiT NanoLuc luciferase subunit; C-Sm-A4galt/B4galt1/B4galt5/B4galt6, human α1,4-galactosyltransferase/β1,4-galactosyltransferase 1/β1,4-galactosyltransferase 5 isoenzyme/β1,4-galactosyltransferase 6 isoenzyme C-terminated fused with the SmBiT NanoLuc luciferase subunit. Statistical significance was established as ∗, p < 0.05; ∗∗p < 0.01; ∗∗∗p < 0.001; and ∗∗∗∗p < 0.0001; ns, not significant.

Additionally, we observed the formation of A4galt-B4galt1 and A4galt-B4galt5 heterodimers in both CHO-Lec2 and HEK293T cells, with no detectable luminescence for the A4galt-B4galt6 pairs (Fig. 1 and 2). The average luminescence measured for A4galt-B4galt1 pairs of high-frequency and mutein enzymes in CHO-Lec2 cells was 90 and 360 times higher than that for controls, respectively (Fig. 2). Similarly, A4galt-B4galt1 heterodimers analyzed in HEK293T cells exhibited robust luminescence signals, approximately 60 times higher for the high-frequency enzyme and approximately 30 times higher for the mutein, compared to the control samples (Fig. 1). The greatest luminescence signals were detected for A4galt-B4galt5 pairs in CHO-Lec2 cells, with approximately 1000 times greater luminescence for the mutein compared to the 300 times higher signal for the high-frequency enzyme, in comparison to controls (Fig. 2). These findings confirmed the ability of human A4galt to form homodimers, as well as heterodimers with B4galt1 and B4galt5, highlighting the varied GSL- and GP-based acceptor specificity.

### Molecular basis of human A4galt dimerization

Our *in vitro* investigations highlighted that human A4galt can form homodimers as well as heterodimers with B4galt1 and B4galt5; however, the mechanism of these PPIs remains elusive. Given the unresolved spatial structure of human A4galt, we employed the AlphaFold-Multimer tool [17] to predict the structures of these complexes. Models for all monomers, including A4galt, B4galt1, and B4galt5, were generated using AlphaFold [18] and are available in the AlphaFold database [36]. The monomer models obtained from the AlphaFold database exhibited high pLDDT scores, indicating a high level of structural reliability, with the exception of the flexible N-terminal fragment. For our modeling endeavors, we considered predictions based on both the full enzyme sequences and those with the N-terminus removed. However, predictions of PPIs using full sequences did not yield reliable outcomes because the N-terminal protein fragment interfered with the formation of intermolecular interfaces in the dimers. Consequently, we focused solely on models with the N-terminus omitted, targeting regions with predictably stable structures for our analysis.

Thus, we attempted to predict the structures of the protein complexes, with the A4galt-B4galt5 heterodimer model emerging as the most accurate prediction (Fig. 3A). This model exhibited the highest pLDDT score, with an average of 92.24. High pLDDT values, particularly those exceeding 90, indicated precise predictions. Furthermore, the high quality of the model was supported by the Predicted Aligned Error (PAE) scores, where the average PAE for residue pairs within a 6 Å contact distance was 3.76 Å (see Fig. SIII). Low PAE values, notably below 5 Å for specific residue pairs, are indicative of a reliable prediction of their spatial relationship, emphasizing the importance of PAE as a key metric for evaluating the structural integrity and precision of protein complexes. Insights into predicted interactions between individual chains were provided by the Mapiya web-server [23], which are elaborated in the Supplementary Data (Fig. 4-5 and SII).

**Fig. 3.**
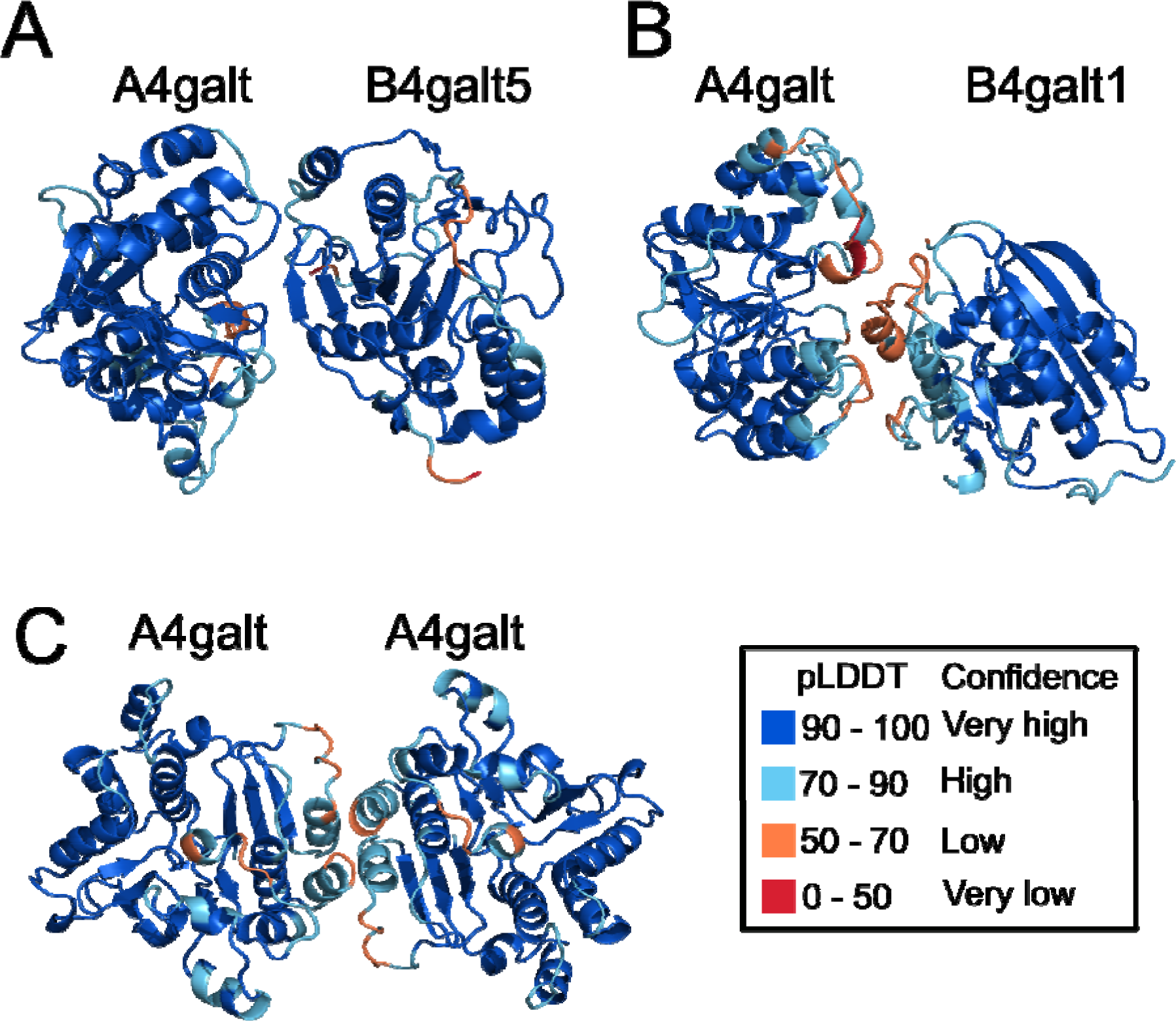
Structure predictions by Alphafold-Multimer: (A) heterodimer A4galt (left) - B4galt1 (right); (B) homodimer A4galt - A4galt; (C) heterodimer A4galt (left) - B4galt5 (right). Coloring based on pLDDT confidence scores.

**Fig. 4.**
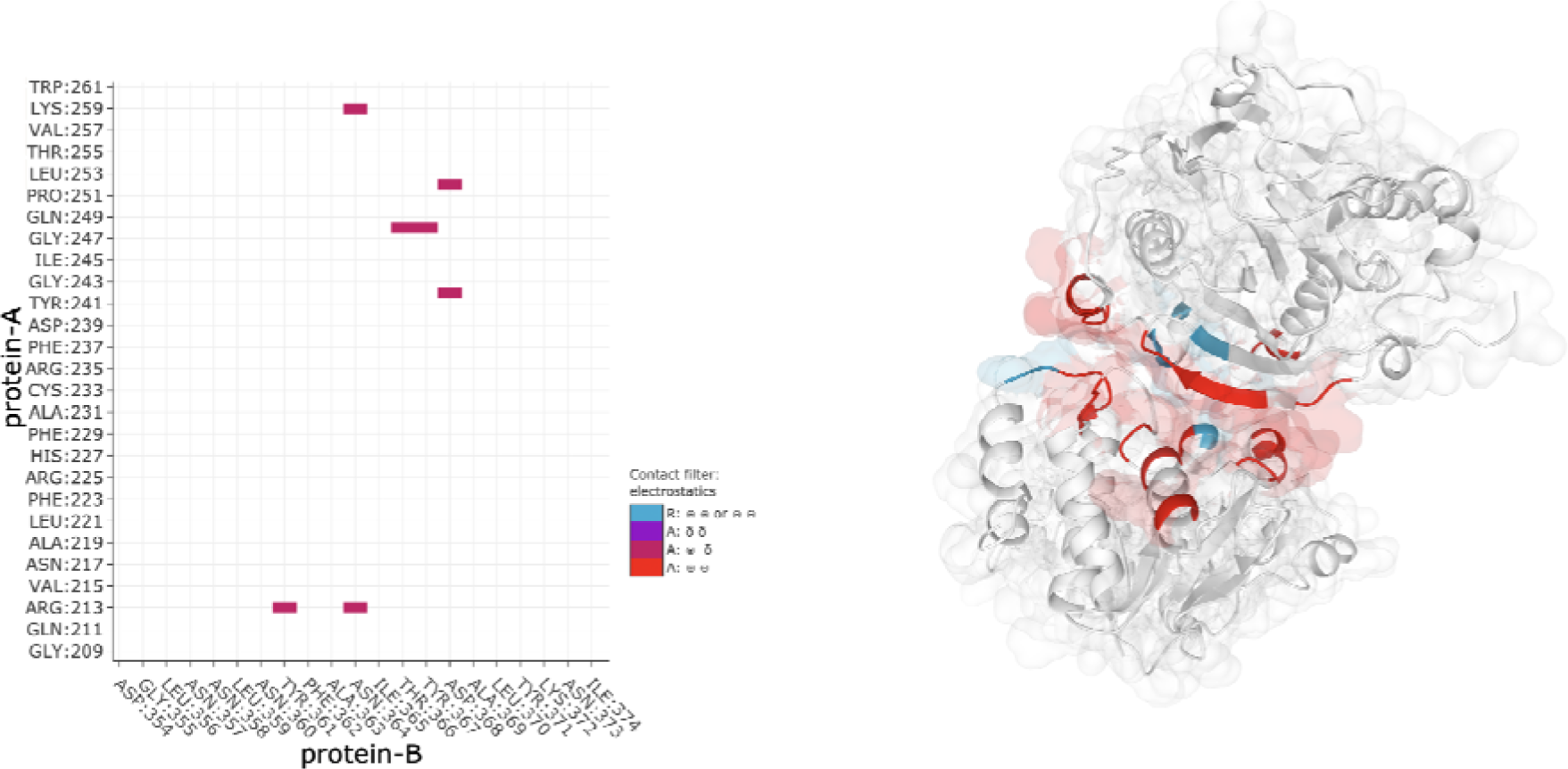
Visualization of pi-stacking interactions between A4galt (bottom, protein-A) and B4galt5 (top, protein-B).

**Fig. 5.**
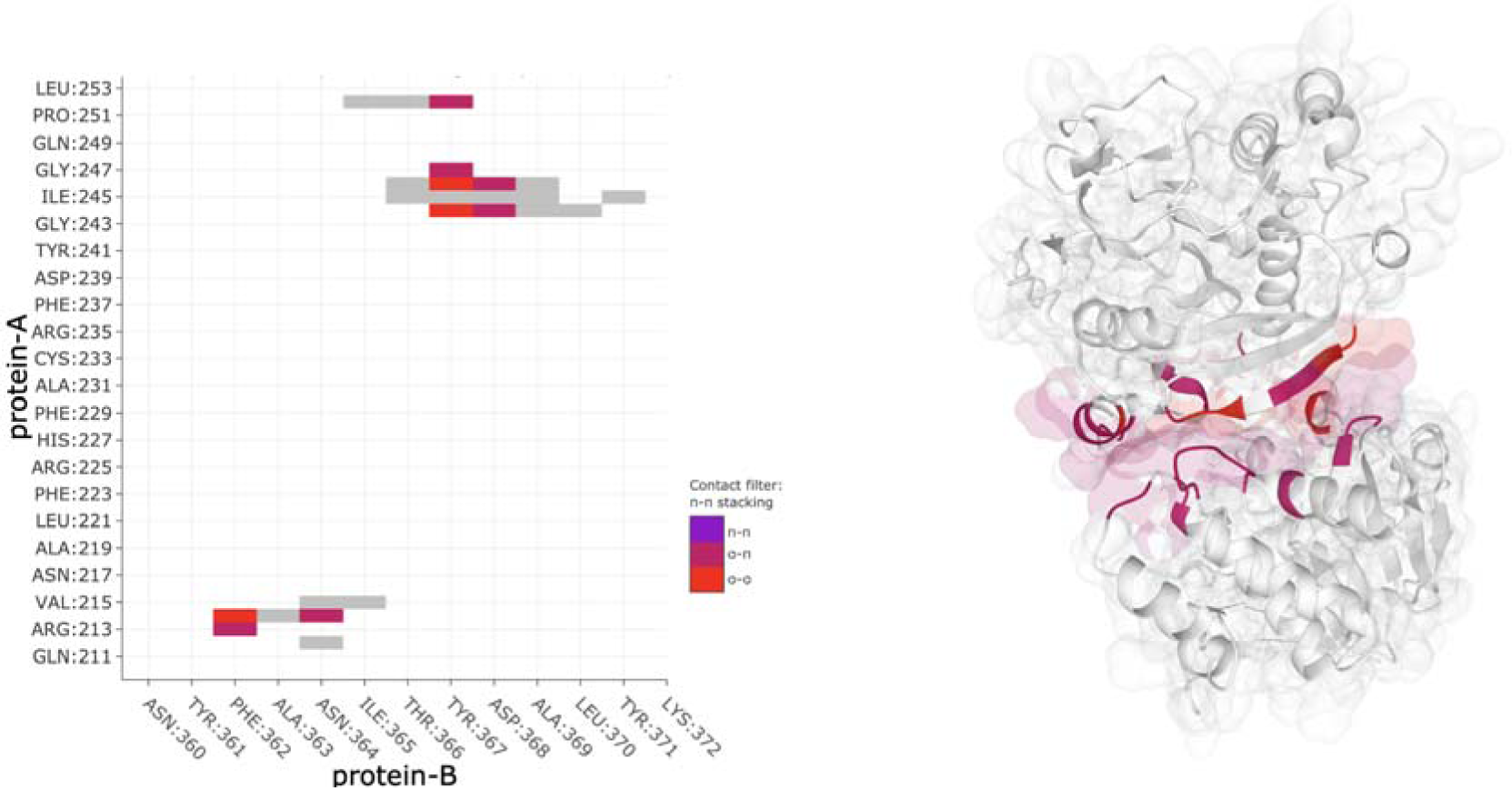
Visualization of electrostatic interactions between A4galt (bottom, protein-A) and B4galt5 (top, protein-B).

**Fig. 6.**
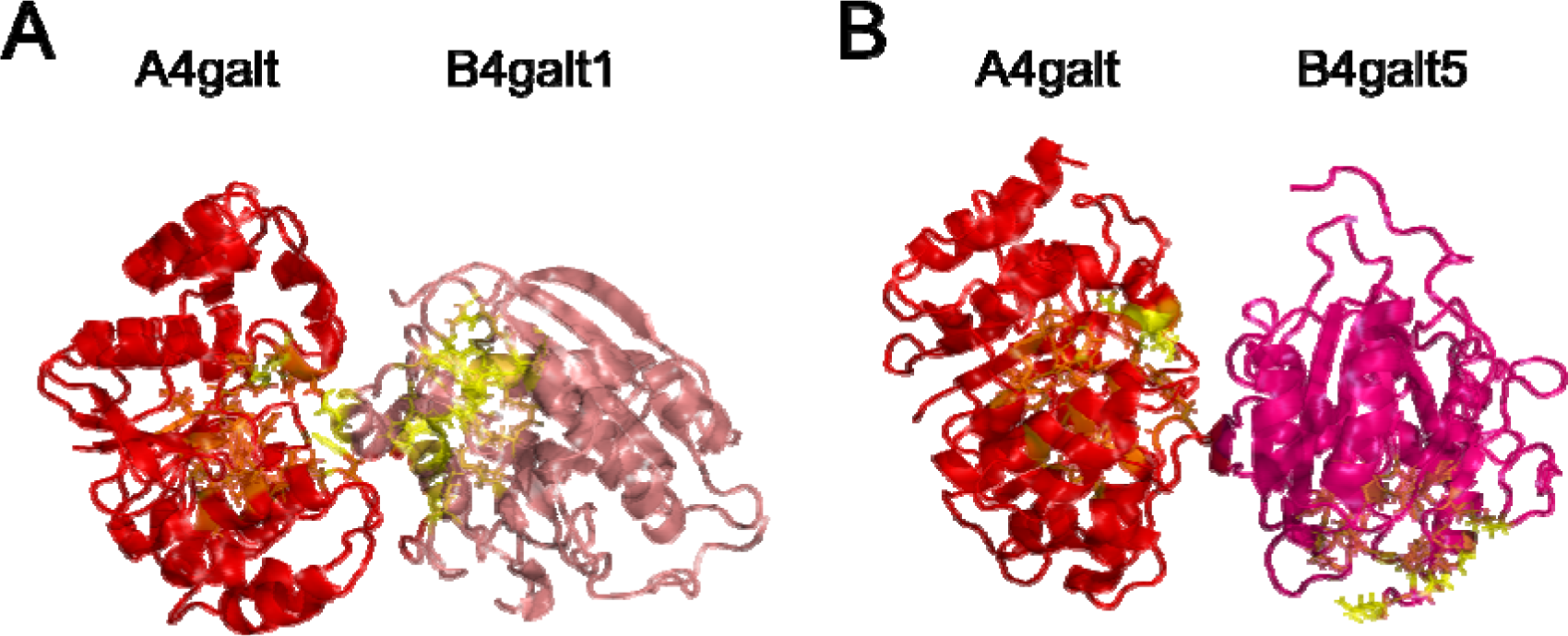
The mutual relationship of active sites (marked by yellow) in heterodimers (A) A4galt (left) - B4galt1 (right); (B) A4galt (left) - B4galt5 (right).

**Fig. 7.**
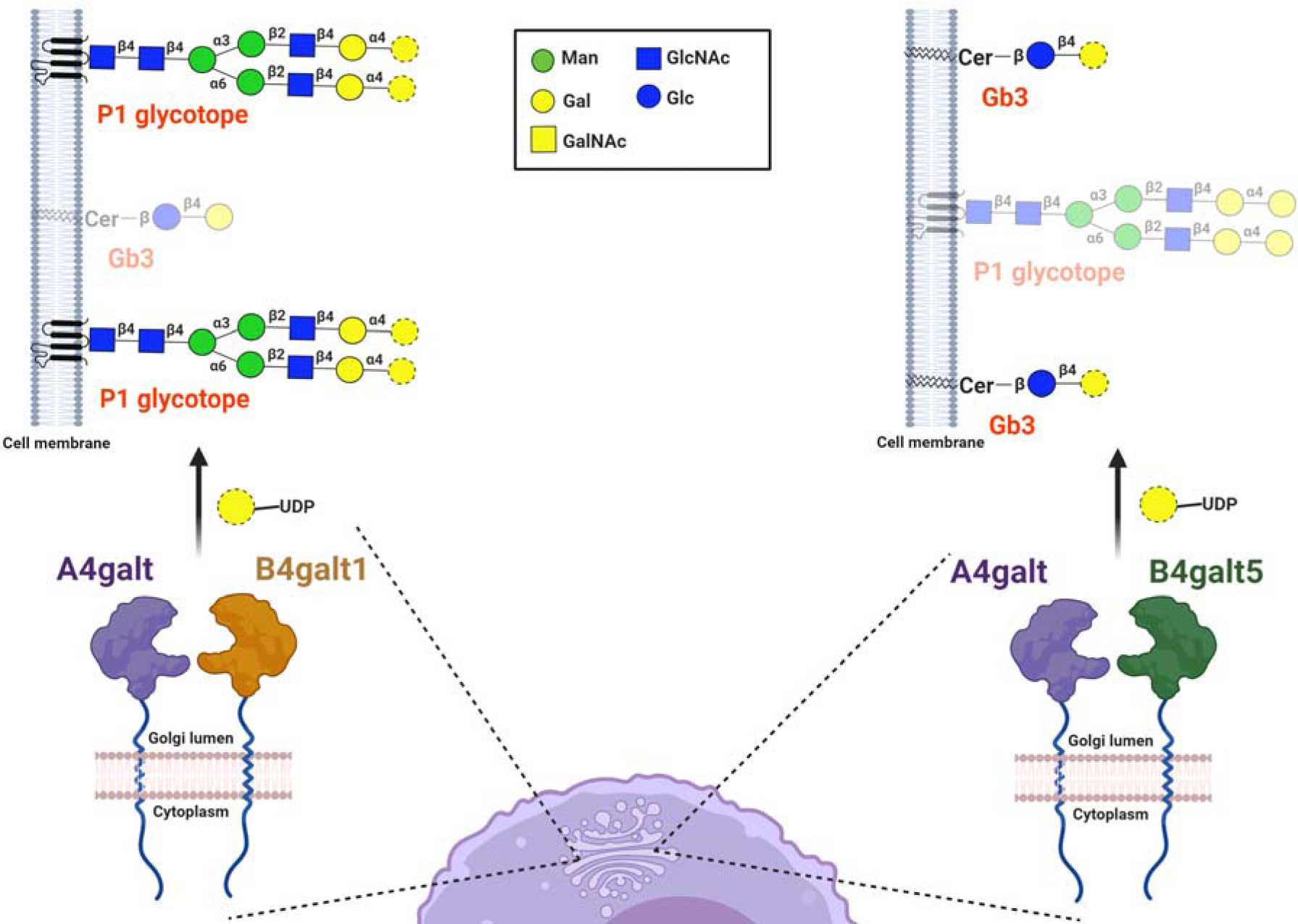
The role of human A4galt heterodimerization with B4galt1 and B4galt5 in determining enzyme acceptor specificity. The formation of the A4galt-B4galt1 heterodimer within the Golgi apparatus is believed to facilitate the recognition of Gal-terminated glycoprotein acceptors by human A4galt, enabling the attachment of a galactose residue to N-glycoproteins [6]. On the other hand, the A4galt-B4galt5 heterodimer seems to be crucial in transferring a galactose residue to Gal-capped glycosphingolipid acceptors, such as lactosylceramide. The coexistence of A4galt-B4galt1 and A4galt-B4galt5 heterodimers within the Golgi apparatus allows these complexes to recognize and modify both GSL- and GP-related substrates, thereby broadening the range of acceptor specificity. The transferred and attached galactose residue by human A4galt was designated with a dashed border.

In addition to the A4galt-B4galt5 model, we also generated models for the homodimer of A4galt-A4galt and a heterodimer of A4galt-B4galt1. However, these complexes exhibited lower pLDDT scores compared to the A4galt-B4galt5 model, particularly within regions where the two monomers interact. Moreover, the high PAE scores for contacting residue pairs in these models suggested diminished confidence in the predicted dimer interface and overall structural configuration (Fig. 3B and C).

Standard AlphaFold-Multimer scores, such as pLLDT and PAE, provide initial insights into the quality of protein structure predictions, but they alone do not guarantee the biological relevance of the interactions. To address this, we have incorporated a machine learning method that enhances prediction accuracy by integrating these scores with a comprehensive set of omics and structural data. This integrated approach is encapsulated in the Structure Prediction and Omics-based Classifier (SPOC), specifically designed to evaluate the validity of predicted protein interactions more effectively [20]. A SPOC score of 0.3 or higher is considered indicative of a high likelihood that the predicted interaction is biologically meaningful and spurious. This threshold was chosen based on its high effectiveness in distinguishing true interactions from computational artifacts, making it a valuable tool for assessing the biological plausibility of our predicted dimer models. In our current study, all three predicted dimer models achieved SPOC scores of 0.3 or above (see Table SIV), suggesting that these models are likely accurate and potentially significant representations of the protein interactions.

To further explore these complexes, we performed coevolution sequence analysis using EVcouplings [21] on the A4galt-B4galt5 heterodimer to identify correlated mutations. The analysis was challenging because of the low number of effective sequences, resulting in a ratio of effective sequences to protein length of 0.47 (the value below 1 suggests the limited reliability of this analysis). We identified only a few residue pairs outside the anticipated interface area, showing only a slight increase in the probability of correlated mutations. A similar coevolution analysis for the A4galt-B4galt1 heterodimer revealed an even smaller number of sequences in the multiple sequence alignment (MSA), highlighting the difficulties in obtaining sufficient confidence in structural predictions for this complex via AlphaFold (Fig. 3B). AlphaFold typically relies on a robust MSA for precision, and the limited number of sequences in MSA poses challenges for accurate predictions.

In addition, we explored the potential active sites of the monomers using the COACH meta-server. We included B4galt1, for which the active site is known [37], to test the reliability of the results. This method proved to be reliable, as it provided consistent predictions across various methods for binding pockets and known substrates associated with these proteins (Fig. SIV-SVI).

Analysis of the mutual arrangement of these active sites, supported by the known structure of the B4galt1 homodimer (PDB ID: 6FWU), suggested that the active sites in the heterodimer A4galt-B4galt1 are proximal enough to facilitate substrate exchange. The AlphaFold-predicted structure showed that most of the interface between the two monomers was composed of residues that formed their respective active centers (Fig. 4A). A similar observation was made for the A4galt-B4galt5 heterodimer, however only the active site residues of A4galt contributed to the interface (Fig. 4B). The proximity between the catalytic centers of A4galt and B4galt1 indicated the potential for substrate transfer.

## Discussion

In this study, we aimed to elucidate the ability of high-frequency and mutein human A4galt to engage in homo- and heterodimerization using the novel NanoBiT technology. Our findings indicate the propensity of human A4galt for homodimerization, however, the functional consequences of this phenomenon for enzyme functions require further evaluation. Homodimerization has been documented in various GTs with distinct acceptor specificities, such as glycoprotein-specific α1,6-fucosyltransferase 8 (Fut8) [38] and glycolipid-specific β1,4-*N*-acetylgalactosaminyltransferase 1 (B4galnt1) [39].

We demonstrated that human A4galt, whether in its high-frequency or mutein form, can form heterodimers with B4galt1 and B4galt5. These heterodimers exhibit distinct acceptor specificity towards GSLs and GPs. Notably, previous research conducted by Takematsu et al. (2011) suggested a potential interaction between human A4galt and B4galt6, which was supported by the shared presence of the GOLPH3 recognition polypeptide motif in these enzymes [16, 19, 40]. To date, many examples of GT heterodimers with varied specificities towards GSLs and GPs have been described. For instance, heterodimers such as glycoprotein-specific β1,4-galactosyltransferase 1/α2,6-sialyltransferase 1 [41] and β1,3-*N*-acetylglucosaminyltransferase 2/β1,3-*N*-acetylglucosaminyltransferase 8 [42], are crucial in N-glycoprotein synthesis. Additionally, glycolipid-specific heterodimers formed by β1,4-*N*-acetylgalactosaminyltransferase 1 with β1,3-galactosyltransferase 4 [43] and α2,8-sialyltransferase 8A with β1,4-*N*-acetylgalactosaminyltransferase 1 [44] play pivotal roles in ganglio-series GSLs synthesis.

Here, we aimed to elucidate the molecular intricacies underlying the A4galt, B4galt1, and B4galt5 interactions using AlphaFold enzyme models. Analysis of the A4galt-B4galt1 heterodimer revealed spatial proximity of the active sites of both enzymes, potentially facilitating mutual substrate exchange. This observation is reminiscent of the mechanism demonstrated by human α2,8-sialyltransferase 3, which forms a homodimer close to the Golgi apparatus membrane [45]. With regards to the A4galt-B4galt5 heterodimer, we proposed an electrostatic bond-mediated enzyme interaction, similar to the mechanism previously described for homo- and heterodimer formation by β1,4-galactosyltransferase 1 and α2,6-sialyltransferase 1 [46]. We assume that both the A4galt-B4galt1 and A4galt-B4galt5 heterocomplexes encourage the transfer of products from B4galt1 and B4galt5 (Gal-terminated N-glycoproteins and glycosphingolipids, respectively) to A4galt. Subsequently, A4galt synthesizes Gal-capped oligosaccharides using two different types of acceptor molecules (GSLs and GPs) [1,6,7].

The role of GT dimerization on acceptor specificity is essential for studying the molecular basis of glycosylation reactions in cells as well as the relationship between GT complexes and the synthesized glycosylated products [14,47–48]. Future studies should focus on elucidating the preferential interactions between A4galt and B4galt5, which seem to be essential for recognizing GSL-based acceptors. More comprehensive studies are required to confirm the existence of direct interactions between specific amino acids in human A4galt and B4galt5 (Fig. SIII). Albeit B4galt5 and B4galt6 isoenzymes reveal very similar specificities, we did not detect the A4galt-B4galt6 heterodimer in either CHO-Lec2 or HEK293F cells. We cannot exclude the competition between B4galt5 and B4galt6 for accessible substrates (a phenomenon described for some B4galts) [18] and/or the mechanistic opportunity to form heterodimers with A4galt by only one isoenzyme; both of which require further investigation, especially unveiling the spatial structure of human A4galt, B4galt5 and B4galt6.

Exploring the molecular underpinnings of the dual acceptor specificity of human A4galt may provide invaluable insights into the pathogenesis of Gb3- and P1 glycotope-related diseases, and contribute to the development of innovative therapeutic strategies. GSL-based Gb3 is a receptor for pathogens, such as uropathogenic *E. coli* strains, zoonotic *Streptococcus nois* and Stxs [49]. Elevated Gb3 levels are associated with Fabry disease, a rare genetic disease caused by α-galactosidase deficiency [50]. Additionally, Gb3 is involved in epithelial-mesenchymal transition, a pivotal process in cancer progression [51]. In contrast, the GP-based P1 glycotope can bind Stx1 and potentially act as a decoy receptor, suggesting its promising therapeutic implications for treating STEC infections in the future [6]. Thus, elucidating the molecular interactions of A4galt may lead to the development of novel targeted therapies, that can mitigate Gb3- and P1-related diseases and offer more effective treatment approaches.

## Author contributions

**Krzysztof Mikolajczyk** (Conceptualization [lead], Data curation [equal], Formal analysis [equal], Funding acquisition [equal], Investigation [equal], Methodology [equal], Project administration [lead], Software [supporting], Supervision [lead], Validation [equal], Visualization [equal], Writing – original draft [equal], Writing – review & editing [equal]), **Karol Wróblewski** (Data curation [equal], Formal analysis [supporting], Investigation [equal], Methodology [equal], Software [lead], Validation [equal], Visualization [equal], Writing – original draft [equal], Writing – review & editing [equal]), **Sebastian Kmiecik** (Conceptualization [supporting], Data curation [supporting], Formal analysis [supporting], Funding acquisition [equal], Investigation [equal], Methodology [equal], Project administration [supporting], Software [supporting], Supervision [supporting], Validation [equal], Visualization [equal], Writing – original draft [equal], Writing – review & editing [equal]).

## Abbreviations

A4galt, α1,4-galactosyltransferase; B4galt, β1,4-galactosyltransferase; Gb3, globotriaosylceramide, Gb3Cer, CD77, P^k^ antigen, ceramide trihexoside, Galα1→4Galβ1→4Glc-Cer; GP, glycoprotein; GL, glycolipid; HUS, hemolytic-uremic syndrome; P1 glycotope, Galα1→4Galβ1→4GlcNAc-R; LacCer, lactosylceramide; NanoBiT, NanoLuc® Binary Technology; PPI, protein-protein interaction; STEC, Shiga toxin-producing *Escherichia coli*; Stx, Shiga toxin; Stx1B, Shiga toxin 1B subunit; Stx2B, Shiga toxin 2B subunit.

## Acknowledgments

The authors are deeply grateful to prof. Marcin Czerwinski for his invaluable insights and thoughtful suggestions throughout the creation of this manuscript. The figures 7 and SI were created with BioRender.com.

