## Supplementary data for "Delving into human α1,4-galactosyltransferase acceptor specificity: the role of enzyme dimerization"

**Contents**

**Table SI.** Nucleotide sequences of the *B4GALTs* and *A4GALT* genes used for cloning into pBiT expression vectors. In underlined **red** marked nucleotide substitution related to mutein enzyme form (rs397514502, c.631C>G, p.Q211E).

**Table SII**. PCR conditions used in the study.

**Table SIII**. pBiT-based expression vectors used in the study. To the pBiT expression vectors were cloned insert genes, including human *A4GALT* (GenBank accession number NM_001318038.3: nucleotides 325-1386 and mutein form with c.631C>G substitution), human *B4GALT1* (GenBank accession number NM_001378495.1: 30-1187), human *B4GALT5* (GenBank accession number NM_004776.4: 189-1355) and human *B4GALT6* (GenBank accession number NM_004775.5: 156-1187), using restriction sites marked in primer sequence as red. The nucleotide sequences of cloned genes in primer sequences were in *italics*. The underlined nucleotides were added to the primer sequences to create 5’-overhangs.

**Table SIV**. The computational assessment scores for predicted protein-protein interactions using Structure Prediction and Omics informed Classifier (SPOC) tool on predictomes.org. Each row represents a different protein pair and includes the SPOC score and 'avg_models' (the fraction of residues that meet certain contact criteria, that are consistently observed across all independently-trained AlphaFold-Multimer models). Additional columns display traditional AlphaFold-Multimer metrics: 'ipTM' - interface predicted Template Modeling score, 'pDOCKQ' - predicted DOCKQ score, 'pLDDT' - predicted local distance difference test, and 'PAE' - predicted alignment error.

**Figure SI.** Schematic representations of the analyzed protein pairs (comprising human B4galt1, B4galt5, B4galt6 and A4galt) to evaluate PPIs using NanoBiT technology.

**Figure SII.** Illustration of PAE score for A4galt-B4galt5 heterodimer. Predicted aligned error (PAE) provides an estimated distance error between pairs of residues.

**Figure SIII.** Potential hydrogen bonds in heterodimer A4galt (bottom, in orange) - B4galt5 (bottom, in green).

**Figure SIV**. Predicted active site for A4galt. Residues forming the active center (marked by yellow): L82, E83, T84, P172, S175, D176, R179, I180, Y190, L191, D192, T193, D194, N217, G218, A219, W246, G247, G250, P251, L254, H315, W317, N318, K319.

**Figure SV**. Predicted active site for B4galt5. Residues forming the active center (marked by yellow): P169, F170, R171, R173, F208, R210, D234, V235, D236, K261, G274, W296, G297, E299, D300, S326, H329, H331, R332.

**Figure SVI**. Predicted active site for B4galt1 (on the left) and the active site in the PDB structure (on the right). Residues forming the active center (marked by yellow): P183, F184, R185, R187, F222, R224, D248, V249, D250, K275, G288, W310, G311, E313, D314, M340, H343, D346, N349.

**Table SI. Nucleotide sequences of the *B4GALTs* and *A4GALT* genes used for cloning into pBiT expression vectors. In underlined red marked nucleotide substitution related to mutein enzyme form (rs397514502, c.631C>G, p.Q211E).**

| **Gene name** | **Nucleotide sequence (5’→3’)** |
| --- | --- |
| *B4GALT1*  (NM_001378495.1: 30-1187) | ATGCCAGGCGCGTCCCTACAGCGGGCCTGCCGCCTGCTCGTGGCCGTCTGCGCTCTGCACCTTGGCGTCACCCTCGTTTACTACCTGGCTGGCCGCGACCTGAGCCGCCTGCCCCAACTGGTCGGAGTCTCCACACCGCTGCAGGGCGGCTCGAACAGTGCCGCCGCCATCGGGCAGTCCTCCGGGGAGCTCCGGACCGGAGGGGCCCGGCCGCCGCCTCCTCTAGGCGCCTCCTCCCAGCCGCGCCCGGGTGGCGACTCCAGCCCAGTCGTGGATTCTGGCCCTGGCCCCGCTAGCAACTTGACCTCGGTCCCAGTGCCCCACACCACCGCACTGTCGCTGCCCGCCTGCCCTGAGGAGTCCCCGCTGCTTGTGGGCCCCATGCTGATTGAGTTTAACATGCCTGTGGACCTGGAGCTCGTGGCAAAGCAGAACCCAAATGTGAAGATGGGCGGCCGCTATGCCCCCAGGGACTGCGTCTCTCCTCACAAGGTGGCCATCATCATTCCATTCCGCAACCGGCAGGAGCACCTCAAGTACTGGCTATATTATTTGCACCCAGTCCTGCAGCGCCAGCAGCTGGACTATGGCATCTATGTTATCAACCAGGCGGGAGACACTATATTCAATCGTGCTAAGCTCCTCAATGTTGGCTTTCAAGAAGCCTTGAAGGACTATGACTACACCTGCTTTGTGTTTAGTGACGTGGACCTCATTCCAATGAATGACCATAATGCGTACAGGTGTTTTTCACAGCCACGGCACATTTCCGTTGCAATGGATAAGTTTGGATTCAGCCTACCTTATGTTCAGTATTTTGGAGGTGTCTCTGCTCTAAGTAAACAACAGTTTCTAACCATCAATGGATTTCCTAATAATTATTGGGGCTGGGGAGGAGAAGATGATGACATTTTTAACAGATTAGTTTTTAGAGGCATGTCTATATCTCGCCCAAATGCTGTGGTCGGGAGGTGTCGCATGATCCGCCACTCAAGAGACAAGAAAAATGAACCCAATCCTCAGAGGTTTGACCGAATTGCACACACAAAGGAGACAATGCTCTCTGATGGTTTGAACTCACTCACCTACCAGGTGCTGGATGTACAGAGATACCCATTGTATACCCAAATCACAGTGGACATCGGGACACCGAGC |
| *B4GALT5* (NM_004776.4: 189-1355) | ATGCGCGCCCGCCGGGGGCTGCTGCGGCTGCCGCGCCGCTCGCTGCTCGCCGCGCTCTTCTTCTTTTCTCTCTCGTCCTCGCTGCTGTACTTCGTCTATGTGGCGCCCGGCATAGTGAACACCTACCTCTTCATGATGCAAGCCCAAGGCATTCTGATCCGGGACAACGTGAGAACAATCGGTGCTCAGGTTTATGAGCAGGTGCTTCGGAGTGCTTATGCCAAGAGGAACAGCAGTGTAAATGACTCAGATTATCCTCTTGACTTGAACCACAGTGAAACCTTCCTGCAAACTACAACATTTCTTCCTGAAGACTTCACCTACTTTGCAAACCATACCTGCCCTGAAAGACTCCCTTCCATGAAGGGCCCAATAGACATAAACATGAGTGAAATTGGAATGGATTACATTCATGAACTCTTCTCCAAAGACCCAACCATCAAGCTCGGAGGTCACTGGAAGCCTTCTGATTGCATGCCTCGGTGGAAGGTGGCGATCCTTATCCCCTTCCGGAACCGCCACGAGCACCTCCCAGTCCTGTTCAGACACCTGCTTCCCATGCTCCAGCGCCAGCGCTTGCAGTTTGCATTTTATGTGGTTGAACAAGTTGGTACCCAACCCTTTAATCGAGCCATGCTTTTCAACGTTGGCTTTCAAGAGGCAATGAAAGACTTGGATTGGGACTGTTTGATTTTTCATGATGTAGATCACATACCGGAAAGTGATCGCAACTATTATGGATGTGGACAGATGCCGAGGCATTTTGCAACCAAATTGGATAAGTATATGTATCTGCTTCCTTATACCGAGTTCTTTGGCGGAGTGAGTGGCTTAACAGTGGAACAATTTCGGAAAATCAATGGCTTTCCTAATGCTTTCTGGGGTTGGGGTGGAGAAGATGACGACCTCTGGAACAGAGTACAGAATGCAGGCTATTCTGTGAGCCGGCCAGAGGGTGACACAGGAAAGTACAAGTCCATTCCTCATCACCATCGAGGAGAAGTCCAGTTTCTTGGAAGGTATGCTCTGCTGAGGAAGTCAAAAGAACGGCAAGGGCTGGATGGCCTCAACAACCTGAACTACTTTGCAAACATCACATACGACGCCTTGTATAAAAACATAACTGTCAACCTGACACCCGAGCTGGCTCAGGTGAACGAGTAC |
| *B4GALT6* (NM_004775.5: 156-1187) | ATGTCTGTGCTCAGGCGGATGATGCGGGTTTCCAATCGCTCTCTCCTCGCCTTCATCTTCTTCTTCTCCCTCTCTTCGTCCTGTCTGTACTTCATCTATGTGGCCCCAGGCATCGCCAACACATATCTCTTTATGGTACAAGCTCGAGGTATAATGTTGAGAGAAAATGTGAAAACAATAGGTCATATGATCAGGCTGTACACAAATAAAAACAGTACGCTCAACGGTACAGATTATCCCGAAGGCAATAATTCAAGTGATTATCTTGTTCAAACAACAACGTATCTCCCGGAAAACTTCACATACTCACCATACCTCCCCTGTCCAGAAAAGCTGCCTTATATGCGAGGATTCCTCAATGTCAATGTAAGCGAAGTCAGTTTTGATGAAATTCATCAACTCTTCTCCAAGGATTTAGATATTGAGCCAGGGGGTCATTGGAGGCCAAAAGACTGTAAACCCAGATGGAAGGTGGCAGTTCTCATTCCTTTCCGTAATCGCCATGAACATCTTCCAATTTTTTTCTTACATCTGATTCCAATGCTCCAGAAGCAGCGGCTGGAATTTGCGTTTTATGTCATTGAACAGACTGGCACACAACCTTTTAACCGTGCGATGCTTTTCAATGTGGGCTTCAAAGAGGCCATGAAAGACAGTGTCTGGGACTGTGTAATCTTCCACGATGTGGATCATCTACCTGAAAATGACCGGAACTATTACGGATGTGGAGAAATGCCACGTCATTTTGCTGCAAAGCTGGATAAATACATGTATATTCTTCCATATAAAGAATTTTTTGGTGGTGTAAGTGGGCTGACAGTGGAACAATTTAGAAAGATCAATGGTTTTCCTAATGCCTTCTGGGGATGGGGAGGAGAAGATGATGACCTTTGGAACAGAGTTCACTATGCTGGATATAATGTAACCAGACCAGAGGGAGACTTAGGAAAATACAAGTCAATTCCTCATCACCATAGAGGTGAAGTCCAGTTTTTAGGACGGTATAAATTACTAAGGTATTCCAAGGAGCGTCAGTACATCGATGGACTGAACAATTTAATATATAGGCCAAAAATACTGGTTGATAGGTTGTATACAAACATATCTGTAAACCTCATGCCAGAGTTAGCTCCAATCGAAGACTAT |
| *A4GALT*  (high frequency, NM_001318038.3: 325-1386) | ATGTCCAAGCCCCCCGACCTCCTGCTGCGGCTGCTCCGGGGCGCCCCAAGGCAGCGGGTCTGCACCCTGTTCATCATCGGCTTCAAGTTCACGTTTTTCGTCTCCATCATGATCTACTGGCACGTTGTGGGAGAGCCCAAGGAGAAAGGGCAGCTCTATAACCTGCCAGCAGAGATCCCCTGCCCCACCTTGACACCCCCCACCCCACCCTCCCACGGCCCCACTCCAGGCAACATCTTCTTCCTGGAGACTTCAGACCGGACCAACCCCAACTTCCTGTTCATGTGCTCGGTGGAGTCGGCCGCCAGAACTCACCCCGAATCCCACGTGCTGGTCCTGATGAAAGGGCTTCCGGGTGGCAACGCCTCTCTGCCCCGGCACCTGGGCATCTCACTTCTGAGCTGCTTCCCGAATGTCCAGATGCTCCCGCTGGACCTGCGGGAGCTGTTCCGGGACACACCCCTGGCCGACTGGTACGCGGCCGTGCAGGGGCGCTGGGAGCCCTACCTGCTGCCCGTGCTCTCCGACGCCTCCAGGATCGCACTCATGTGGAAGTTCGGCGGCATCTACCTGGACACGGACTTCATTGTTCTCAAGAACCTGCGGAACCTGACCAACGTGCTGGGCACCCAGTCCCGCTACGTCCTCAACGGCGCGTTCCTGGCCTTCGAGCGCCGGCACGAGTTCATGGCGCTGTGCATGCGGGACTTCGTGGACCACTACAACGGCTGGATCTGGGGTCACCAGGGCCCGCAGCTGCTCACGCGGGTCTTCAAGAAGTGGTGTTCCATCCGCAGCCTGGCCGAGAGCCGCGCCTGCCGCGGCGTCACCACCCTGCCCCCTGAGGCCTTCTACCCCATCCCCTGGCAGGACTGGAAGAAGTACTTTGAGGACATCAACCCCGAGGAGCTGCCGCGGCTGCTCAGTGCCACCTATGCTGTCCACGTGTGGAACAAGAAGAGCCAGGGCACGCGGTTCGAGGCCACGTCCAGGGCACTGCTGGCCCAGCTGCATGCCCGCTACTGCCCCACGACGCACGAGGCCATGAAAATGTACTTGTGA |
| *A4GALT*  (mutein, NM_001318038.3: 325-1386 with c.631C>G substitution, rs397514502) | ATGTCCAAGCCCCCCGACCTCCTGCTGCGGCTGCTCCGGGGCGCCCCAAGGCAGCGGGTCTGCACCCTGTTCATCATCGGCTTCAAGTTCACGTTTTTCGTCTCCATCATGATCTACTGGCACGTTGTGGGAGAGCCCAAGGAGAAAGGGCAGCTCTATAACCTGCCAGCAGAGATCCCCTGCCCCACCTTGACACCCCCCACCCCACCCTCCCACGGCCCCACTCCAGGCAACATCTTCTTCCTGGAGACTTCAGACCGGACCAACCCCAACTTCCTGTTCATGTGCTCGGTGGAGTCGGCCGCCAGAACTCACCCCGAATCCCACGTGCTGGTCCTGATGAAAGGGCTTCCGGGTGGCAACGCCTCTCTGCCCCGGCACCTGGGCATCTCACTTCTGAGCTGCTTCCCGAATGTCCAGATGCTCCCGCTGGACCTGCGGGAGCTGTTCCGGGACACACCCCTGGCCGACTGGTACGCGGCCGTGCAGGGGCGCTGGGAGCCCTACCTGCTGCCCGTGCTCTCCGACGCCTCCAGGATCGCACTCATGTGGAAGTTCGGCGGCATCTACCTGGACACGGACTTCATTGTTCTCAAGAACCTGCGGAACCTGACCAACGTGCTGGGCACC**G**AGTCCCGCTACGTCCTCAACGGCGCGTTCCTGGCCTTCGAGCGCCGGCACGAGTTCATGGCGCTGTGCATGCGGGACTTCGTGGACCACTACAACGGCTGGATCTGGGGTCACCAGGGCCCGCAGCTGCTCACGCGGGTCTTCAAGAAGTGGTGTTCCATCCGCAGCCTGGCCGAGAGCCGCGCCTGCCGCGGCGTCACCACCCTGCCCCCTGAGGCCTTCTACCCCATCCCCTGGCAGGACTGGAAGAAGTACTTTGAGGACATCAACCCCGAGGAGCTGCCGCGGCTGCTCAGTGCCACCTATGCTGTCCACGTGTGGAACAAGAAGAGCCAGGGCACGCGGTTCGAGGCCACGTCCAGGGCACTGCTGGCCCAGCTGCATGCCCGCTACTGCCCCACGACGCACGAGGCCATGAAAATGTACTTGTGA |

**Table SII**. PCR conditions used in the study.

| **PCR step** | **Parameters** | | |
| --- | --- | --- | --- |
|  | **Temp [°C]** | **Time [s]** | **Cycle** |
| **Initial denaturation** | 94 | 180 | 1 |
| **Denaturation** | 94 | 30 | 30 |
| **Annealing** | 58 | 20 |  |
| **Extension** | 72 | 30 |  |
| **Final extension** | 72 | 300 | 1 |

**Table SIII**. pBiT-based expression vectors used in the study. To the pBiT expression vectors were cloned insert genes, including human *A4GALT* (GenBank accession number NM_001318038.3: nucleotides 325-1386 and mutein form with c.631C>G substitution), human *B4GALT1* (GenBank accession number NM_001378495.1: 30-1187), human *B4GALT5* (GenBank accession number NM_004776.4: 189-1355) and human *B4GALT6* (GenBank accession number NM_004775.5: 156-1187), using restriction sites marked in primer sequence as red. The nucleotide sequences of cloned genes in primer sequences were in *italics*. The underlined nucleotides were added to the primer sequences to create 5’-overhangs.

| **Construct** | **Insert** | **Tag** | **Original plasmid** | **Primer sequence [5’ – 3’]** | **Restriction site** |
| --- | --- | --- | --- | --- | --- |
| N-S-B4G1 | Human B4galt1 | SmBiT or  LgBiT at the N-terminus | pBiT2.1-N[TK/SmBiT]  or  pBiT1.1-N[TK/LgBiT] | FP: AAAACTCGAGA*ATGCCAGGCGCGTCCCTAC* | *XhoI* |
|  |  |  |  | RP: AAAAGAATTC*CTAGCTCGGTGTCCCGATGTCC* | *EcoRI* |
| C-S-B4G1 | Human B4galt1 | SmBiT or  LgBiT at the C-terminus | pBiT2.1-C[TK/SmBiT] or  pBiT1.1-C[TK/LgBiT] | FP: GCCGCAGAATTCA *ATGCCAGGCGCGTCCCTAC* | *EcoRI* |
|  |  |  |  | RP: AAAACTCGAGCC*GCTCGGTGTCCCGATGTCC* | *XhoI* |
| N-L-B4G5 | Human B4galt5 | SmBiT or  LgBiT at the N-terminus | pBiT2.1-N[TK/SmBiT]  or  pBiT1.1-N[TK/LgBiT] | FP: AAAACTCGAGA*ATGCGCGCCCGCCGGG* | *XhoI* |
|  |  |  |  | RP: ATATCGGAATTC*CTAGTACTCGTTCACCTGAGCCAG* | *EcoRI* |
| C-S-B4G5 | Human B4galt5 | SmBiT or  LgBiT at the C-terminus | pBiT2.1-C[TK/SmBiT] or  pBiT1.1-C[TK/LgBiT] | FP: ATATCGGAATTCA *ATGCGCGCCCGCCGGG* | *EcoRI* |
|  |  |  |  | RP: AAAACTCGAGCC*GTACTCGTTCACCTGAGCCAG* | *XhoI* |
| N-L-B4G6 | Human B4galt6 | SmBiT or  LgBiT at the N-terminus | pBiT2.1-N[TK/SmBiT]  or  pBiT1.1-N[TK/LgBiT] | FP: AAAACTCGAGA *ATGTCTGTGCTCAGGCGGATG* | *XhoI* |
|  |  |  |  | RP: ATATCGGAATTC*CTAATAGTCTTCGATTGGAGCTAACT* | *EcoRI* |
| C-S-B4G6 | Human B4galt6 | SmBiT or  LgBiT at the C-terminus | pBiT2.1-C[TK/SmBiT] or  pBiT1.1-C[TK/LgBiT] | FP: ATATCGGAATTCA*ATGTCTGTGCTCAGGCGGATG* | *EcoRI* |
|  |  |  |  | RP: AAAACTCGAGCC*ATAGTCTTCGATTGGAGCTAACT* | *XhoI* |
| N-S-A4G | Human A4galt | SmBiT or  LgBiT at the N-terminus | pBiT2.1-N[TK/SmBiT]  or  pBiT1.1-N[TK/LgBiT] | FP: AAAA CTCGAGA*ATGTCCAAGCCCCCCGAC* | *XhoI* |
|  |  |  |  | RP: AAAA GAATTC *TCACAAGTACATTTTCATGGC* | *EcoRI* |
| C-S-A4G | Human A4galt | SmBiT or  LgBiT at the C-terminus | pBiT2.1-C[TK/SmBiT] or  pBiT1.1-C[TK/LgBiT] | FP: AAAAGAATTC*ATGTCCAAGCCCCCCGAC* | *EcoRI* |
|  |  |  |  | RP: AAAA CTCGAGCC*CAAGTACATTTTCATGGCCTC* | *XhoI* |

**Table SII**. The computational assessment scores for predicted protein-protein interactions using Structure Prediction and Omics informed Classifier (SPOC) tool on predictomes.org. Each row represents a different protein pair and includes the SPOC score and 'avg_models' (the fraction of residues that meet certain contact criteria, that are consistently observed across all independently-trained AlphaFold-Multimer models). Additional columns display traditional AlphaFold-Multimer metrics: 'ipTM' - interface predicted Template Modeling score, 'pDOCKQ' - predicted DockQ score, 'pLDDT' - predicted local distance difference test, and 'PAE' - predicted alignment error.

| **Pair name** | **SPOC score** | **avg_models** | **ipTM** | **pDockQ** | **pLDDT** | **PAE** |
| --- | --- | --- | --- | --- | --- | --- |
| A4galt:A4galt | 0.46 | 0.621 | 0.47 | 0.15 | 71.3 | 10.4 |
| A4galt:B4galt5 | 0.30 | 0.333 | 0.82 | 0.22 | 90.2 | 2.8 |
| A4galt:B4galt1 | 0.30 | 0.333 | 0.27 | 0.15 | 67.9 | 14.9 |

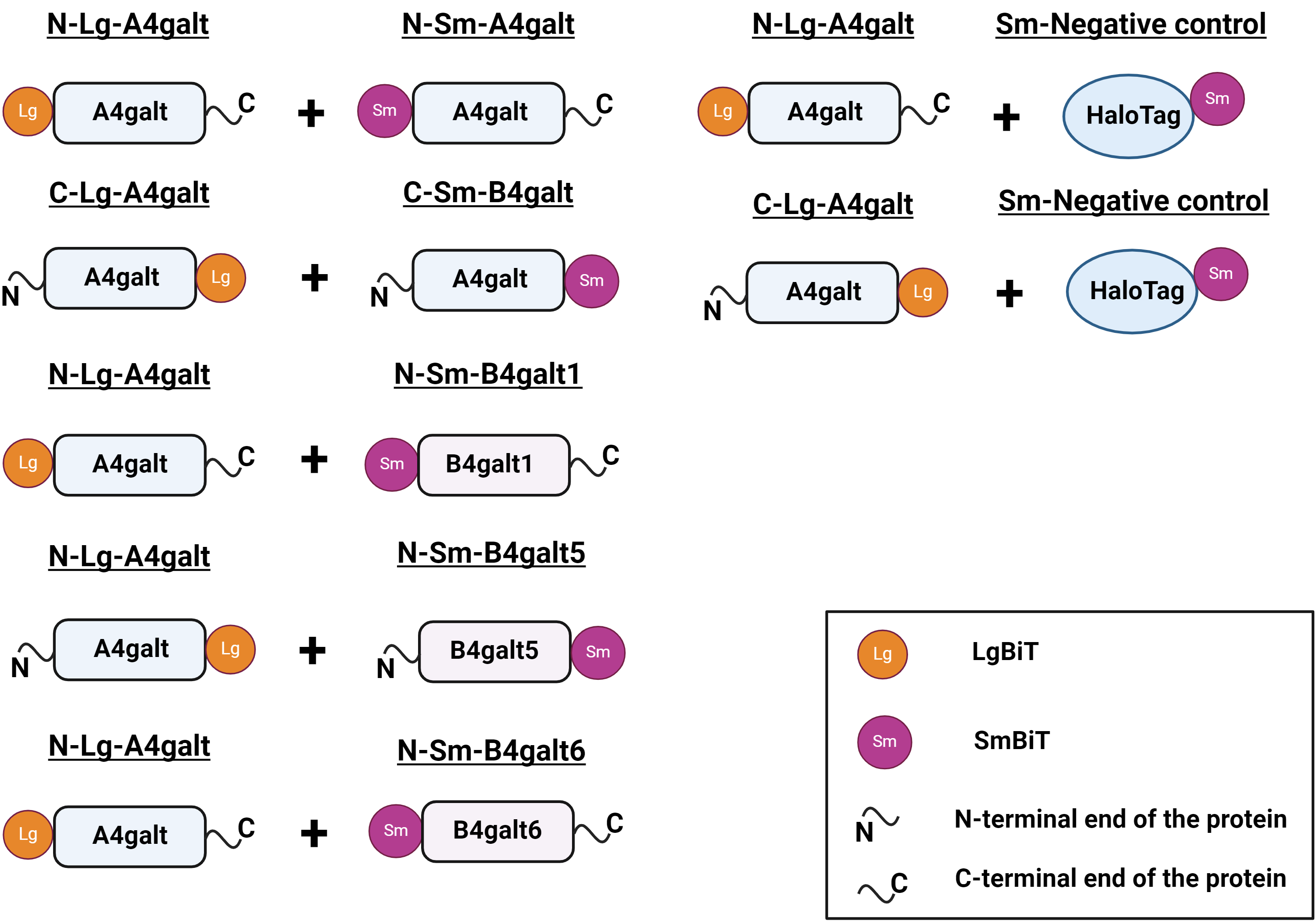

**Fig. SI.** Schematic representations of the analyzed protein pairs (comprising human B4galt1, B4galt5, B4galt6 and A4galt) to evaluate PPIs using NanoBiT technology.

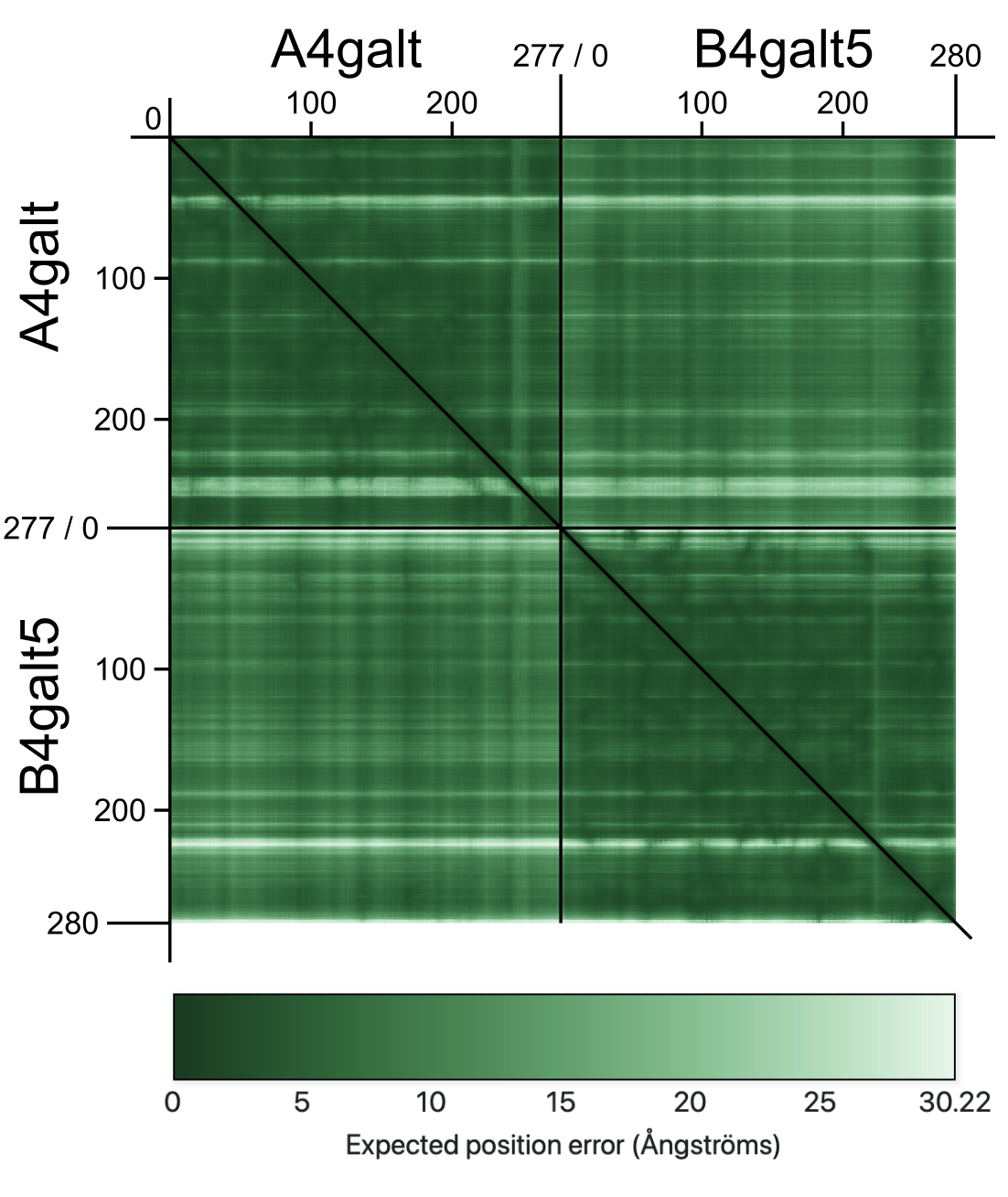

**Fig. SII.** Illustration of PAE score for A4galt-B4galt5 heterodimer. Predicted aligned error (PAE) provides an estimated distance error between pairs of residues.

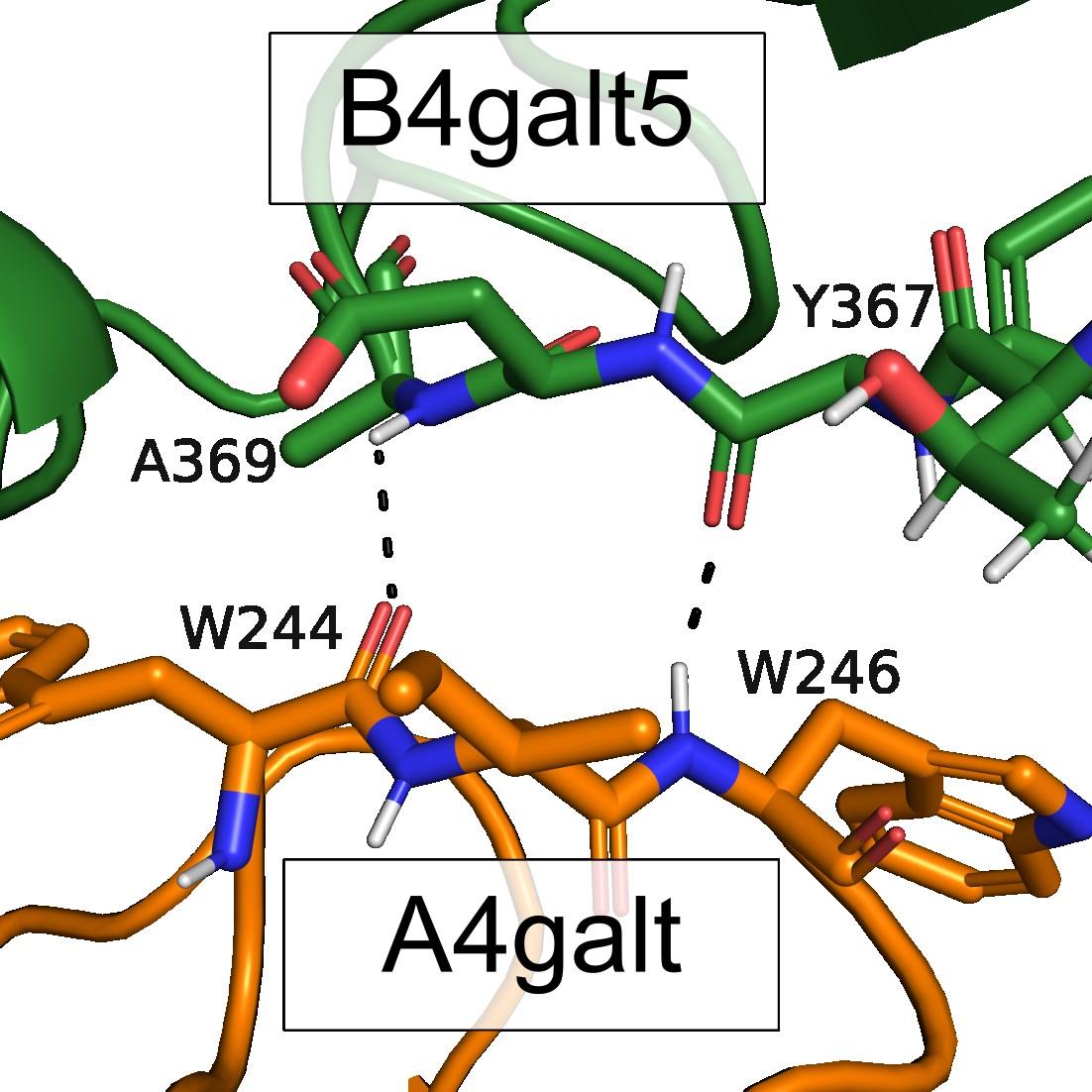

**Fig. SIII.** Potential hydrogen bonds in heterodimer A4galt (bottom, in orange) - B4galt5 (bottom, in green).

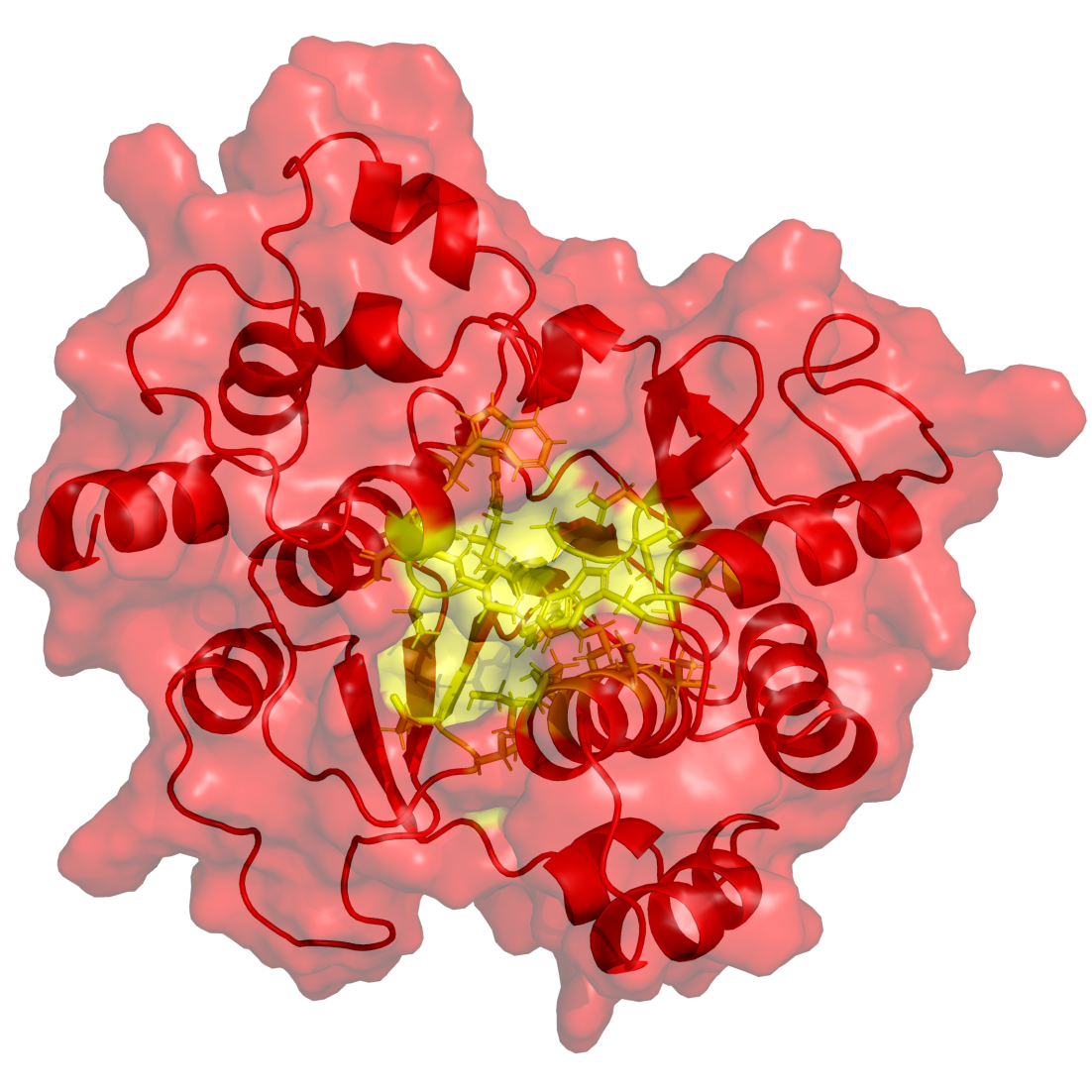

**Fig. SIV.** Predicted active site for A4galt. Residues forming the active center (marked by yellow): L82, E83, T84, P172, S175, D176, R179, I180, Y190, L191, D192, T193, D194, N217, G218, A219, W246, G247, G250, P251, L254, H315, W317, N318, K319.

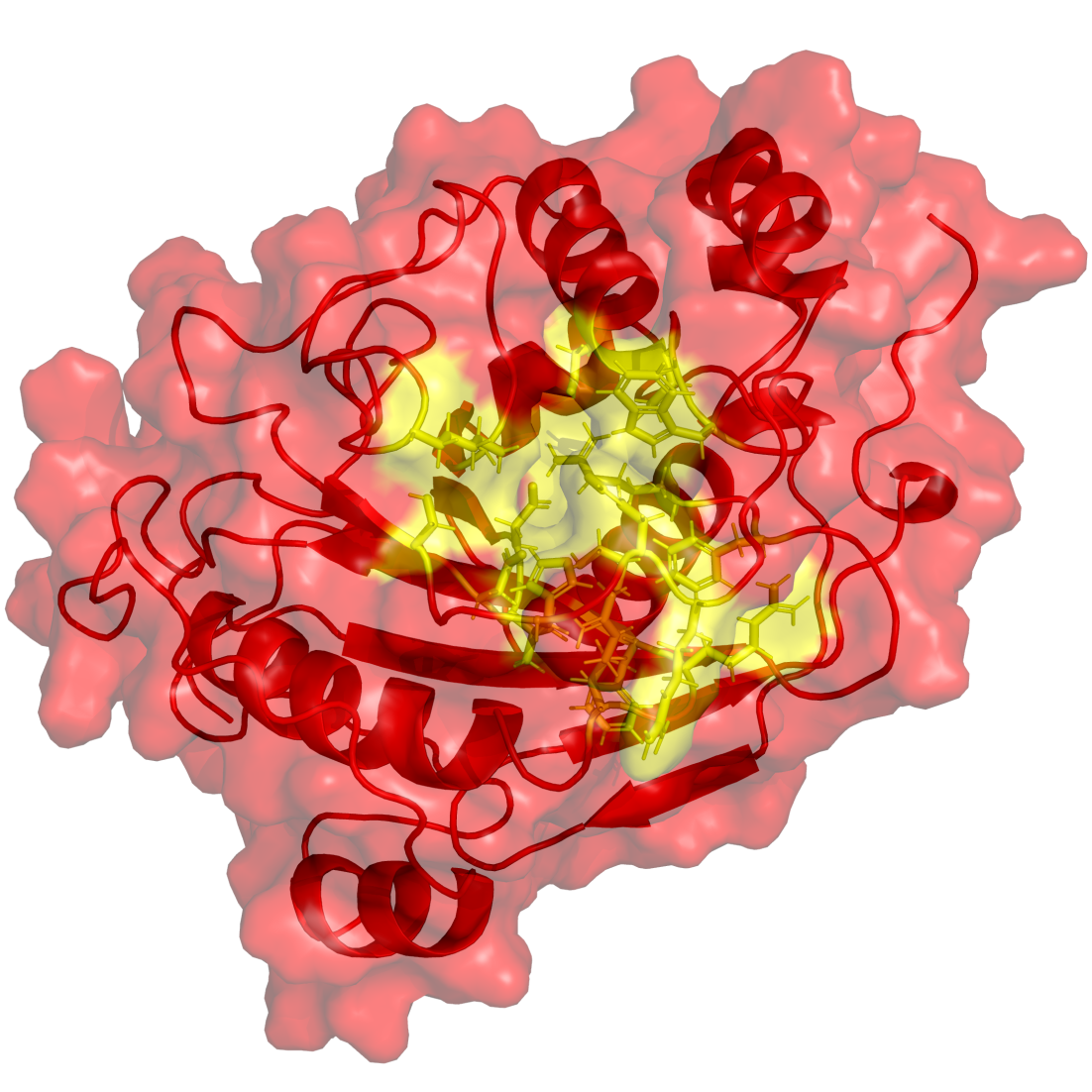

**Fig. SV.** Predicted active site for B4galt5. Residues forming the active center (marked by yellow): P169, F170, R171, R173, F208, R210, D234, V235, D236, K261, G274, W296, G297, E299, D300, S326, H329, H331, R332.

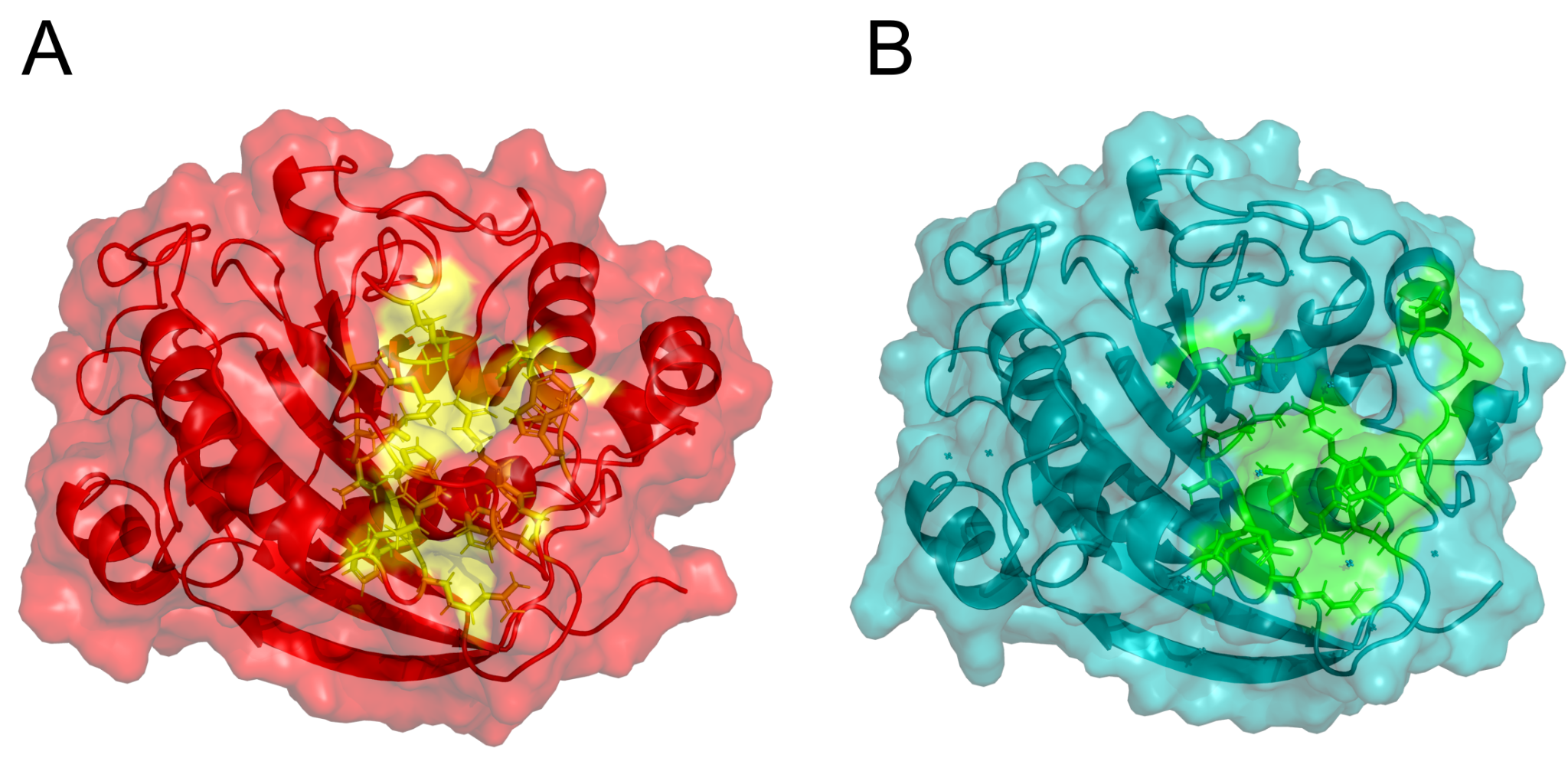

**Fig. SVI.** Predicted active site for B4galt1 (A) and the active site in the PDB structure (B). Residues forming the active center: P183, F184, R185, R187, F222, R224, D248, V249, D250, K275, G288, W310, G311, E313, D314, M340, H343, D346, N349.
